# Binding partners regulate unfolding of myosin VI to activate the molecular motor

**DOI:** 10.1101/2020.05.10.079236

**Authors:** Ália dos Santos, Natalia Fili, Yukti Hari-Gupta, Rosemarie E. Gough, Lin Wang, Marisa Martin-Fernandez, Jesse Aaron, Eric Wait, Teng-Leong Chew, Christopher P. Toseland

## Abstract

Myosin VI is the only minus-end actin motor and is coupled to various cellular processes ranging from endocytosis to transcription. This multi-potent nature is achieved through alternative isoform splicing and interactions with a network of binding partners. There is a complex interplay between isoforms and binding partners to regulate myosin VI. Here, we have compared the regulation of two myosin VI splice isoforms by two different binding partners. By combining biochemical and single-molecule approaches, we propose that myosin VI regulation follows a generic mechanism, independently of the spliced isoform and the binding partner involved. We describe how myosin VI adopts an autoinhibited backfolded state which is released by binding partners. This unfolding activates the motor, enhances actin binding and can subsequently trigger dimerization. We have further expanded our study by using single molecule imaging to investigate the impact of binding partners upon myosin VI molecular organisation and dynamics.

## INTRODUCTION

Myosins are actin-based molecular motors which perform vital roles in numerous of cellular processes [1]. Myosin VI (MVI) is associated with several cellular functions, ranging from endocytosis to transcription [2–9]. MVI is unique, in that it is the only member of the myosin family with the ability to move towards the minus end of actin filaments [10] and its functional diversity relies on its association with various binding partners [11, 12].

MVI is comprised of a motor domain, followed by a neck region consisting of a unique insert, which confers the reverse directionality, and an IQ domain (Figure 1A). Both of these domains bind calmodulin. The N-terminal tail domain contains three structural domains: a three-helix-bundle (amino acids 835-916)[13], a single-alpha-helix (amino acids 942-978), followed by a short coiled-coil [14]. The C-terminal tail domain consists of the globular cargo binding domain (CBD 1060-end). In addition, two regions within the tail can be alternatively spliced resulting in a 31-residue insertion (large-insert, LI) proximal to the CBD, and/or an 9-residue insertion within the CBD (small-insert, SI). This leads to four splice isoforms, the non-insert (NI), SI, LI and LI+SI [15], each with distinct intracellular distributions and functions [15, 16]. For example, the MVI-NI isoform is able to enter the nucleus, whereas the MVI-LI is confined to the cell periphery [4].

**Figure 1:**
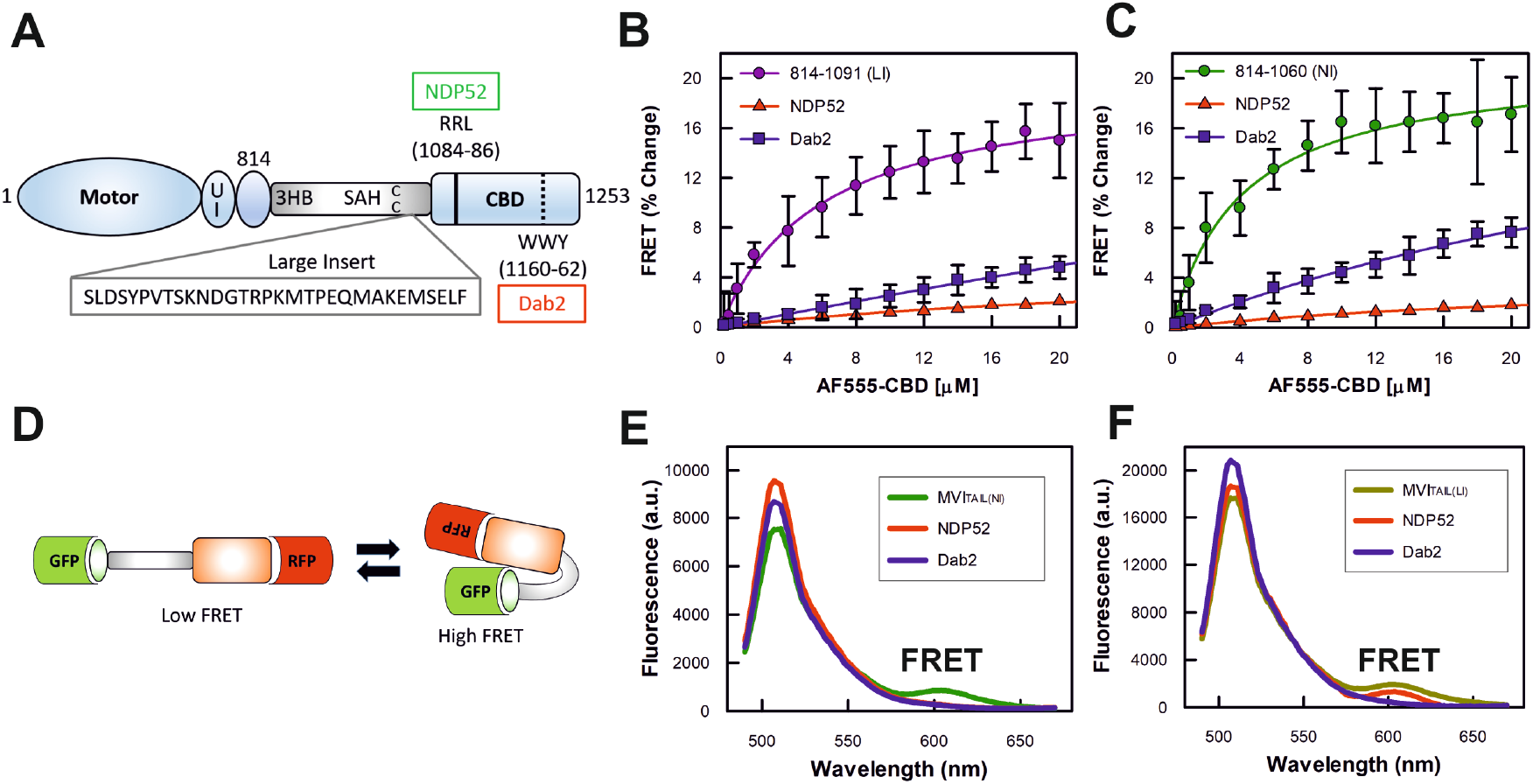
Backfolding of the myosin VI tail. (A) Cartoon depiction of the key regions of MVI, as discussed in the text. UI Unique Insert; 3HB Three Helix Bundle; SAH Stable Alpha Helix; CC Coiled-coil; CBD Cargo Binding Domain. This highlights position of the large insert (LI), along with NDP52 and Dab2 binding sites. (B) FRET titration of Alexa555-CBD against 1 μM of FITC-MVI_TAIL(LI)_, large insert (LI) containing MVI_TAIL_ (residues 814-1091), in the presence of NDP52 or tDab2, at 10 μM. (C) FRET titration of Alexa555-CBD against 1 μM of FITC-MVI_TAIL(NI)_ non-insert (NI) MVI_TAIL_ (residues 814-1060), in the presence of NDP52 or tDab2, at 10 μM. All titration data fitting was performed as described in Methods (error bars represent SEM from three independent experiments). (D) Schematic representation of FRET assay to measure backfolding of the MVI_TAIL_. (E) Representative fluorescence spectra of 1 μM GFP-MVI_TAIL(NI)_-RFP +/- 5 μM NDP52, or 20 μM tDab2. (F) Representative fluorescence spectra of 1 μM GFP-MVI_TAIL(LI)_-RFP(right) +/- 5 μM NDP52, or tDab2.

The CBD domain enables interactions with several binding partners which control the intracellular localisation and function of MVI [1]. This diverse list of partners includes disabled-2 (Dab2) and nuclear dot protein 52 (NDP52). These partners predominantly bind to one of the two established motifs within the CBD of MVI, the WWY and RRL, respectively [8, 17, 18]. However, interactions occur over wider regions. Binding partner selectivity is driven by isoform splicing, whereby the MVI-LI encodes an alpha helix which occludes the RRL motif [19]. This prevents partners, such as NDP52, from interacting with the protein, and therefore interactions for this isoform can be driven by the WWY motif. In contrast, in the MVI-NI isoform, both motifs are available for binding. However, the RRL site can display higher affinity for partners over the WWY motif, in order to select for those interactions [6]. We have previously revealed that MVI-NI can adopt a back-folded conformation, in which the CBD is brought into close proximity to the motor domain [4, 20, 21]. NDP52 then interacts through the RRL binding motif, which leads to unfolding and subsequent dimerization of MVI through an internal dimerization site.

Several questions remain unanswered: It is unknown if the structurally distinct MVI-LI isoform is regulated in the same manner, or whether the WWY site can trigger the same structural rearrangements in MVI. Do binding partners enable unfolding of MVI or stabilise an open conformation? How do micromolar protein affinities regulate MVI in the cell? Lastly, how does the release of backfolding regulate motor activity?

To this end, we have now expanded our studies to assess whether this mechanism applies generally to MVI, independent of partner and isoform preference. We have also further explored the molecular basis for this mechanism, and investigated the cellular organisation and dynamics with respect to binding partners. Overall, we present a detailed generic model governing the activation of MVI from an inactive back-folded state to an active motor capable of actin binding and dimerization.

## RESULTS

### Myosin VI back-folding is independent of isoform

We have previously shown that the MVI-NI isoform is back-folded *in vitro* and in cells [4]. To address whether back-folding is a generic feature of MVI, independent of the isoform, we investigated the conformation of MVI-LI. We utilised a previously employed FRET-based assay [4, 6, 22] by titrating Alexa555-CBD against FITC-MVI_814-1091_ (containing the LI) to assess whether there is an interaction between the N- and C-terminal tail domains, as would occur in a back-folded state. A significant concentration-dependent change in FRET was measured, indicating that the two domains are in close proximity (Figure 1B). The same was observed for the MVI-NI isoform MVI_814-1060_ (Figure 1C), consistent with previous results [4]. The calculated equilibrium dissociation constants (*K*_d_) corresponding to these data were 5.98 (+/0.58) μM and 4.8 (+/- 0.61) μM for MVI-LI- and MVI-NI, respectively. These are relatively low micromolar affinities, suggesting the interactions are likely to be dynamic.

### Binding partners regulate back-folded myosin VI

We have previously shown [4] that unfolding of the MVI-NI tail is directly driven by NDP52 following its interaction with the RRL motif and not through Calcium-Calmodulin interactions, as proposed by a different study [20], and we have previously addressed this discrepancy [4]. To address whether regulation of backfolding by binding partners is a generic mechanism, we followed a similar approach for both MVI-LI and MVI-NI isoforms, but now focusing upon the WWY partner of MVI, Dab2. Due to the instability of recombinant full-length Dab2, we used a recombinant C-terminal truncation of the protein (residues 649-770), which contains the MVI binding site [8], as performed previously [6]. This truncation of Dab2 will be referred to as tDab2 throughout the manuscript. To assess whether Dab2 can regulate back-folding, the FRET assay between the CBD and the MVI_814-1091_ (LI) or the MVI_814-1060_ (NI) tail was repeated following pre-incubation of the CBD with an excess of tDab2. As with NDP52, tDab2 sequestered the CBD, preventing the interaction between the two domains (Figures 1B and 1C), suggesting that Dab2 is also able to disrupt the intramolecular back-folding in both isoforms. However, tDab2 was not as efficient as NDP52, which is consistent with its weaker affinity for MVI [6].

To further confirm the direct effect of binding partners upon MVI unfolding, we used the FRET-based MVI tail conformation reporter, which we have previously developed [4]. Our reporter was based on a GFP-RFP FRET pair with the MVI_TAIL(NI)_ or MVI_TAIL(LI)_, placed in the middle (Figure 1D). As shown by the fluorescent spectra, a high FRET population was observed with both of these reporters, supporting the conclusion that the tails of both isoforms have the ability to back-fold (Figure 1E and 1F). The addition of 5 μM NDP52 to the MVI_TAIL(NI)_ FRET reporter, resulted in loss of the high FRET population. Similarly, 5 μM tDab2 was able to deplete the FRET population of the MVI_TAIL(LI)_. However, 20 μM of tDab2 were required to induce an equivalent effect on MVI_TAIL(NI)_, given its low affinity for this tail [6]. Also as expected, NDP52 induced little, if any, loss of the high FRET population of the MVI_TAIL(LI)_ reporter, given that the RRL site is masked by the LI [19].

Altogether, these data demonstrate that the intramolecular backfolding of MVI is not isoform specific, but rather an intrinsic feature of the protein. Moreover, binding partners interacting at either motif can regulate this backfolding by triggering the unfolding step.

### Binding partners associate to back-folded myosin VI and trigger unfolding

Stopped-flow transient kinetics were then employed to further explore the role of binding partner interactions during the unfolding process. In particular, we endeavoured to determine whether binding partners first interact with the backfolded MVI triggering its unfolding, or whether they bind to spontaneously unfolded MVI stabilising the conformation. Given that both MVI isoforms showed the same response for either binding partner tested, further experiments only focused upon the NI isoform and NDP52.

First, FITC-MVI_TAIL(NI)_ and AF555-NDP52 were used as a FRET pair to report upon the interaction. 1 μM FITC-MVI_TAIL(NI)_ was mixed with excess AF555-NDP52 under pseudo-first order conditions (Figure 2A). The fluorescence traces were characterised by two phases: an increase in FRET signal, followed by a partial decrease (Figure 2B). The first phase was fitted to a signal exponential function (Figure 2C) and the observed rate constant was found to be dependent on the concentration of NDP52 (Figure 2D). We extracted an association rate constant of 1.72 μM^-1^ s^-1^ and dissociation rate constant of 3.3 s^-1^, giving a *K*_d_ of 1.9 μM. This is consistent with the Equilibrium Dissociation constant previously derived from titrations [6]. The second phase in all three traces was also fitted to a single exponential function (Figure 2C), however the derived rate constants (average 2.1 s^-1^) were independent of NDP52 concentration (Figure 2D). This indicates that the second phase corresponds to a first order process, such as a conformation change. We therefore propose that these biphasic traces directly report upon a two-step process: first, binding of the partner onto backfolded MVI, corresponding to the initial increase in FRET, and second, the subsequent unfolding of the myosin, corresponding to the consequent decrease in FRET due to a greater distance between the donor-acceptor dyes (Figure 2E). This suggests that direct binding of the unfolded MVI is unlikely to occur.

**Figure 2:**
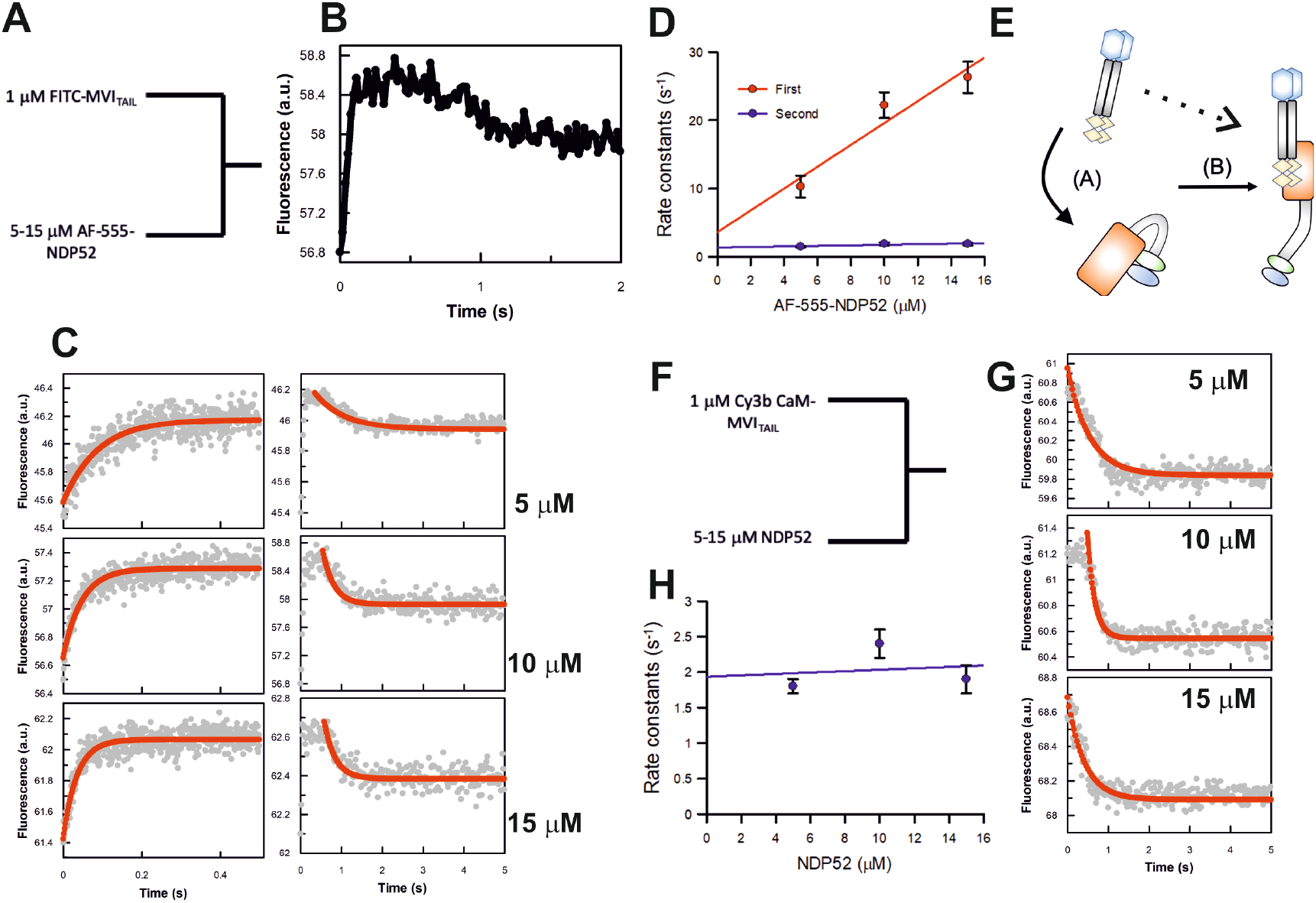
NDP52 association kinetics with the myosin VI tail. (A) Schematic of the experiments for rapid mixing of the FITC labelled MVI_TAIL_ and Alexa-Fluor555 labelled NDP52. Experiments were performed as described in the methods. (B) Representative stopped-flow fluorescence trace depicting both changes in FRET signal, as described in the text. (C) Representative stopped-flow fluorescence traces and exponential fitting to both transitions when the FItC-MVI_TAIL_ is mixed with the stated concentrations of NDP52. (D) The individual traces were fitted to single exponentials and the dependence of the rate constants on concentration was then fitted to a straight line, as shown. The points shown are averages of at least 3 measurements, where error bars represent SEM. The fit for the first phase gives a slope of 1.72 μM^−1^ s^−1^ and an intercept of 3.3 s^−1^. The second phase is independent of NDP52 concentration and gives an average rate constant of 2.1 s^-1^. (E) Cartoon depicting the two processes reported by the stopped-flow experiments. Step A represents NDP52 binding to the MVI_TAIL_, which is dependent upon NDP52 concentration. Step B represents the subsequent unfolding of MVI with a rate constant of 2.1 s^-1^, which is independent of NDP52 concentration. The dotted line represents an alternative model where NDP52 would bind to spontaneously unfolding MVI_TAIL_. (F) Schematic of the experiments for rapid mixing of 1 μM Cy3B-calmodulin-bound MVI_TAIL(NI)_ (pre-mix molar ratio 2:1) and unlabelled NDP52. Experiments were performed as described in the methods. (G) Representative stopped-flow fluorescence traces and exponential fitting to the fluorescence decrease when Cy3B-calmodulin MVI_TAIL(NI)_ is mixed with the stated concentrations of NDP52. (H) The individual traces were fitted to single exponentials and the dependence of the rate constants on NDP52 concentration was then fitted to a straight line, as shown. The points shown are averages of at least 3 measurements, where error bars represent SEM. The rate constants are independent of NDP52 concentration, with an average value of 2 s^-1^.

To further understand this partner-induced conformation change, we used fluorescently labelled calmodulin bound to the MVI_TAIL(NI)_ as an environmental reporter. This approach would enable the probe to report upon either the partner binding step or the MVI unfolding step, or both processes (Figure 2F and G). Cy3B-calmodulin was pre-mixed with MVI_TAIL(NI)_ in a 2:1 molar excess. 1 μM Cy3B-calmodulin MVI_TAIL(NI)_ was then mixed against an excess of non-fluorescent NDP52, under pseudo-first order conditions. A single exponential decrease in fluorescence was observed for all concentrations tested (Figure 2F), suggesting a single step process, which does not arise solely due to Cy3B-calmodulin reacting to NDP52 (Supplementary Figure 1). Interestingly, the derived rate constants (2 s^-1^) were independent of NDP52 concentration (Figure 2H) and were identical to the rate of constants of the second exponential phase observed in the AF555-NDP52 experiments (Figure 2D). We therefore propose that the Cy3B-calmodulin probe reports upon unfolding. This is not unanticipated because unfolding would lead to the largest local environmental change for the dye, as the partner itself binds to the CBD not the neck region. These observations are consistent with the partners not binding to a spontaneously unfolded MVI but rather to its backfolded conformation, which then triggers the unfolding of the protein (Figure 2E).

### Myosin VI dimerization is an intrinsic property

Unfolding of the NI isoform subsequently exposes dimerization sites, leading to protein oligomerization [4], similar to the LI isoform with binding partners [6]. There is a debate as to whether MVI dimerizes intrinsically or through a binding partner mediator [4, 13, 23–25]. We have previously reported that a tail region ahead of the CBD (Figure 3A) of MVI can dimerize independently of binding partners, but we also suggested that this region is blocked until interactions with binding partners occur [4].

**Figure 3:**
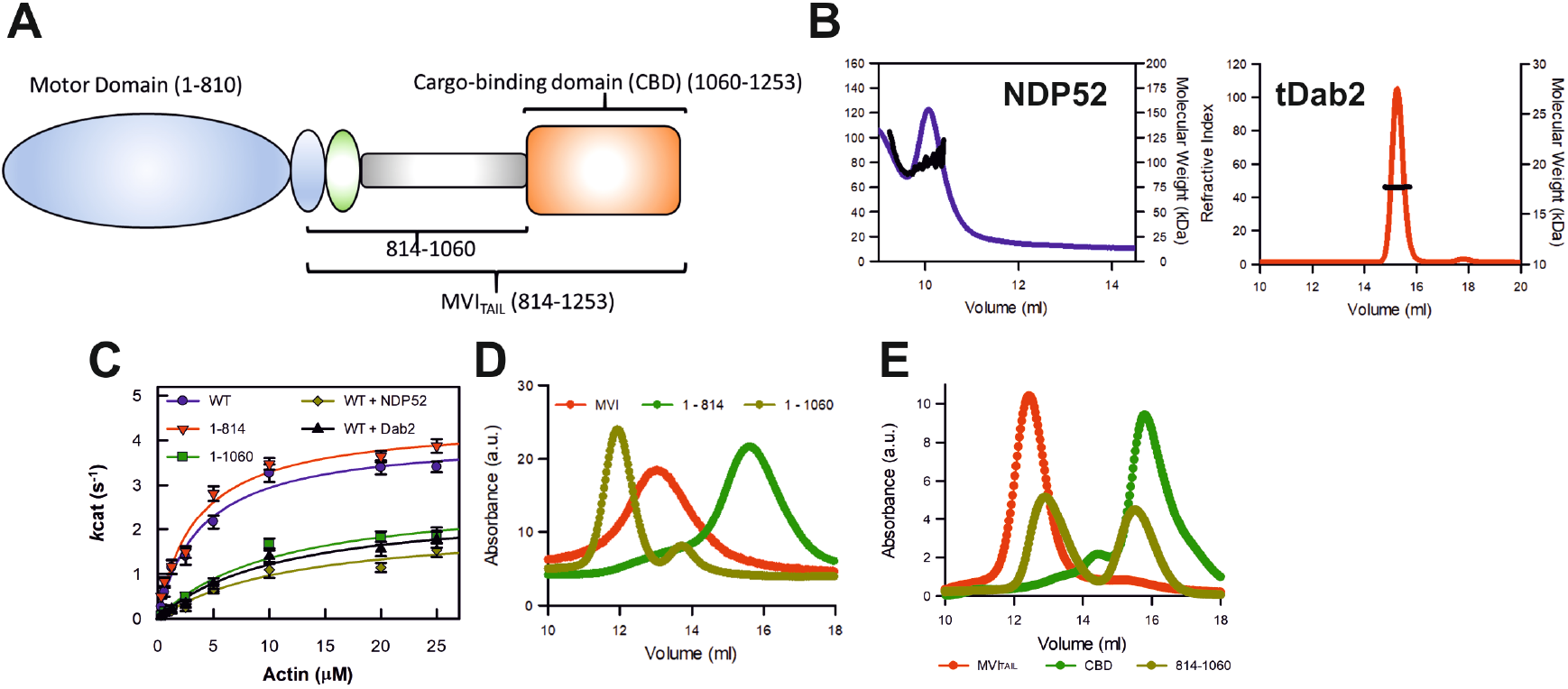
Dimerization of myosin VI. (A) Cartoon depiction of the key domains of MVI which are used in the ATPase and size-exclusion chromatography measurements. (B) Representative SECMALS traces for NDP52 and tDab2 giving molecular weights of 102 kDa and 17 kDa, respectively. This corresponds to dimeric NDP52 and monomeric tDab2. (C) Michaelis-Menten plot displaying steady-state actin-activated ATPase activity for the MVI constructs. Error bars represent SEM from three-independent experiments. (D) Representative SEC traces for 1 mg/ml of the MVI constructs. (E) Representative SEC traces for 1 mg/ml of the MVI_TAIL_ constructs.

NDP52 is a dimeric protein (Figure 3B) and therefore capable of dimerizing MVI with one CBD bound to each monomer. However, tDab2, which also has been previously shown to be able to dimerize MVI [6], is monomeric (Figure 3B). This finding reinforces our previous conclusion that dimerization is an intrinsic property of MVI. This allowed us to estimate that the complex would consist of one binding partner (monomer or dimer) per MVI molecule. For example, we would expect 4 NDP52 molecules (or 2 NDP52 dimers) in complex with 2 MVI proteins.

To explore this hypothesis, we performed ATPase measurements on two MVI constructs, namely MVI_1-814_ and MVI_1-1060_, with the latter lacking the CBD and therefore having the proposed dimerization region exposed (Figure 3C). The ATPase rates for MVI_1-814_ (*k*_cat_ 4.4 s^-1^) were similar to full-length MVI (*k*_cat_ 4.1 s^-1^), while MVI_1-1060_ displayed a lower ATPase rate (*k*_cat_ 2.79 s^-1^), as expected for a dimeric protein. This occurs due to molecular gating, whereby the ATPase activity of the individual motors is coordinated so that only a single motor turns over ATP at any given moment [26]. This rate was also similar to those measured for full-length MVI in the presence of dimeric NDP52 (*k*_cat_ 2.11 s^-1^) and monomeric tDab2 (*k*_cat_ 2.55 s^-1^). Overall, these ATPase measurements show that dimerization is a feature of MVI and not due to binding partner crosslinking. In addition, size-exclusion chromatography strongly supported the formation of MVI_1-1060_ dimers. Full-length MVI eluted as a single peak around 13 ml, whereas MVI_1-814_ eluted at 15.6 ml (Figure 3D), consistent with this construct being smaller in size. However, MVI_1-1060_ eluted earlier than both full length MVI and MVI_1-814_ constructs, suggesting it has adopted a distinct structure which supports its dimeric nature observed in the ATPase measurements. Experiments were also performed with MVI and NDP52, however the proteins separated during the chromatography, so a complex could not be resolved. Finally, size-exclusion chromatography also showed that MVI_814-1060_ elutes in two equal peaks, indicating monomer and dimer species. In contrast, MVI_TAIL(NI)_ and CBD elute as largely single species (Figure 3E), further supporting our finding that dimerization occurs within amino acids 814-1060. Taken together, the ATPase and size-exclusion chromatography data further support the presence of a dimerization site between amino acids 814-1060, consistent with our previous FRET experiments [4, 6].

Lastly, in order to follow the binding partner induced dimerization in real-time, we performed a FRET-based stopped flow assay. 1 μM FITC-MVI_TAIL(NI)_ and 1 μM AF555-MVI_TAIL(NI)_ were pre-mixed, before being mixed in the stopped-flow with an excess of NDP52, under pseudo-first order conditions (Figure 4A). The fluorescence trace revealed a single exponential increase in FRET signal, with an observed rate constant of 2.2 s^-1^ (Figure 4B, C). This was independent of NDP52 concentration (Figure 4C), but importantly, it was similar to the unfolding kinetics of the MVI_TAIL(NI)_ (Figure 2C and 2G. Altogether, we propose that dimerization is an intrinsic property of MVI and is a rapid process that occurs once unfolding has been triggered by the interaction with the binding partners, exposing the otherwise masked dimerization site (Figure 4D).

**Figure 4:**
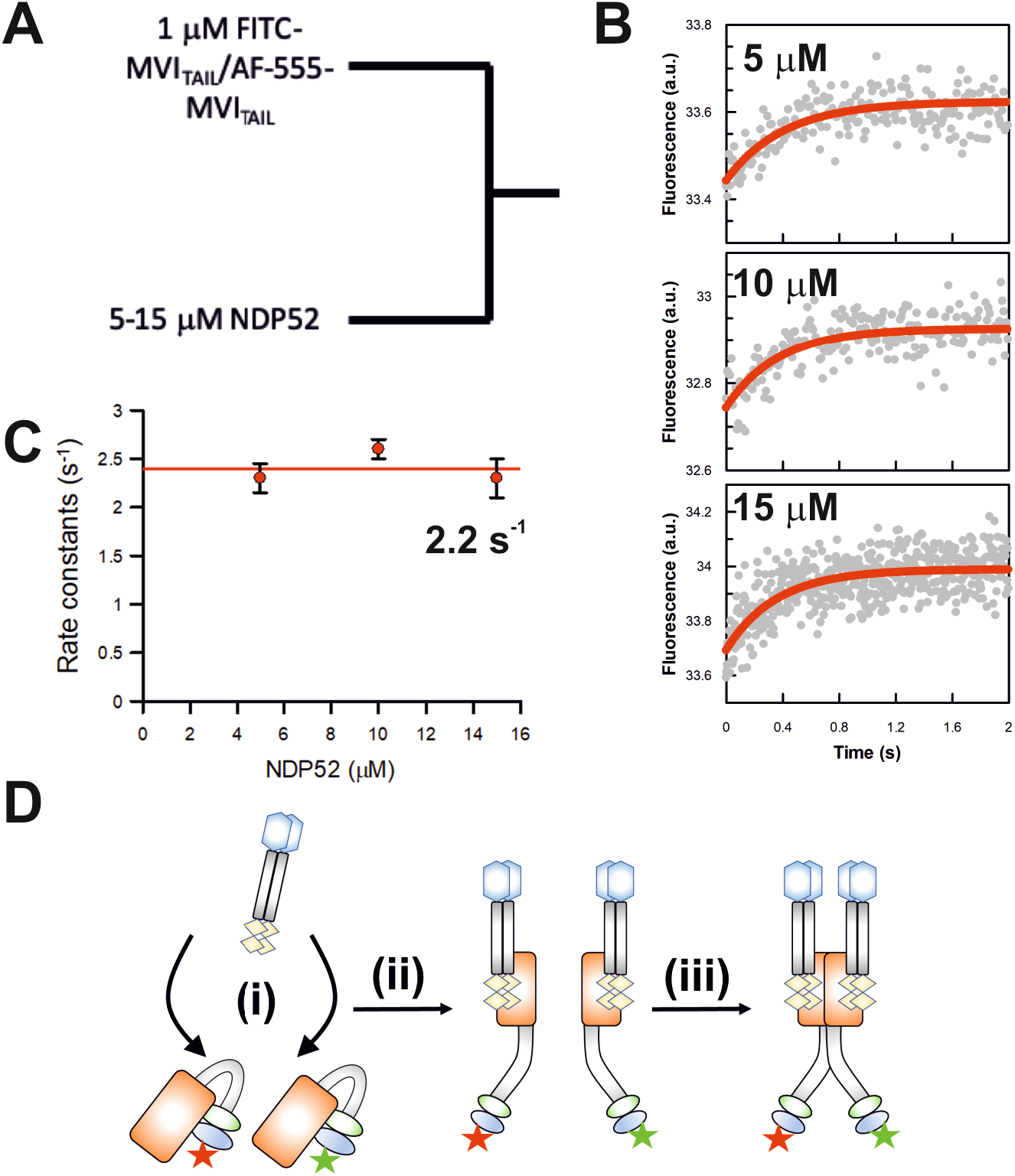
Binding partner driven dimerization of Myosin VI. (A) Schematic of the experiments for rapid mixing of premixed FITC-MVI_TAIL_ and Alexa-Fluor555--MVI_TAIL_ with unlabelled NDP52. Experiments were performed as described in the methods. (B) Representative stopped-flow fluorescence traces and exponential fitting to the fluorescence increase, when the two labelled pools of MVI_TAIL_ are mixed with the stated concentrations of NDP52. (C) The individual traces were fitted to single exponentials and the dependence of the rate constants on concentration was then fitted to a straight line, as shown. The points shown are averages of at least 3 measurements and error bars represent SEM. The rate constants are independent of NDP52 concentration, with an average value of 2.2 s^-1^. (D) Cartoon depicting binding partner driven dimerization based upon the experiments in Figure 2, 3 and 4. Both pools of labelled MVI_TAIL_ are folded. NDP52 binds to either Tail (i) and then triggers their unfolding (ii). This then enables dimerization of the MVI_TAIL_ domains (iii).

### Clustering of myosin VI generates high local densities within the cell

We have explored the molecular mechanism underpinning MVI unfolding and dimerization. However, the affinities determined here and previously [6] are in the low micromolar range. Whilst it is possible that these interactions are part of large multi-valent complexes which enhances the overall complex affinities, the biochemical constants defined here suggest that binding partner interactions and subsequent dimerization would be rare events. We therefore focused upon investigating the *in cellulo* spatial organisation of MVI and NDP52 using super resolution imaging – Stochastic Optical Reconstruction Microscopy (STORM) to investigate cellular interactions.

Widefield imaging against endogenous NDP52 and MVI showed that both proteins are distributed throughout HeLa cells (Figure 5A). STORM imaging of both proteins resolved NDP52 and MVI clusters (Figure 5B). To determine whether this distribution is indeed clustered or random (Figure 5C), we performed cluster analysis using the linearized form of Ripley’s K function [27] *L(r)-r*, where r is the radius. A plot of *L(r)-r* versus *r* gives a value of zero for a random distribution (blue spots), but deviates from zero, due to molecular clustering (Figure 5C). This analysis showed that both NDP52 and MVI assemble into clusters, rather than being randomly distributed. To further understand this clustering behaviour, we used the Clus-DoC software [27], which allows to quantify the spatial distribution of a protein by generating cluster maps (Figure 5D). In this way, we calculated that, in the cytoplasm, 65 % and 70 %of MVI and NDP52 molecules, respectively, are in clusters (Figure 5E). Overexpression of Halo-MVI reveals an increase in the mean number percentage of MVI molecules in clusters (Figure 5E), which suggests clustering is concentration dependent. Interestingly, mutation of the RRL binding sites leads to a 50% reduction in the percentage of molecules in clusters (Figure 5E), which links binding partner interactions to molecular clustering. Lastly, treatment with small molecule inhibitor, TIP, results in a significant decrease in the percentage of molecules in a cluster (Figure 5E). Moreover, TIP leads to a significant decrease in the total number of clusters and number of molecules per cluster (Supplementary Figure 2), suggesting that molecular clustering requires MVI activity.

**Figure 5:**
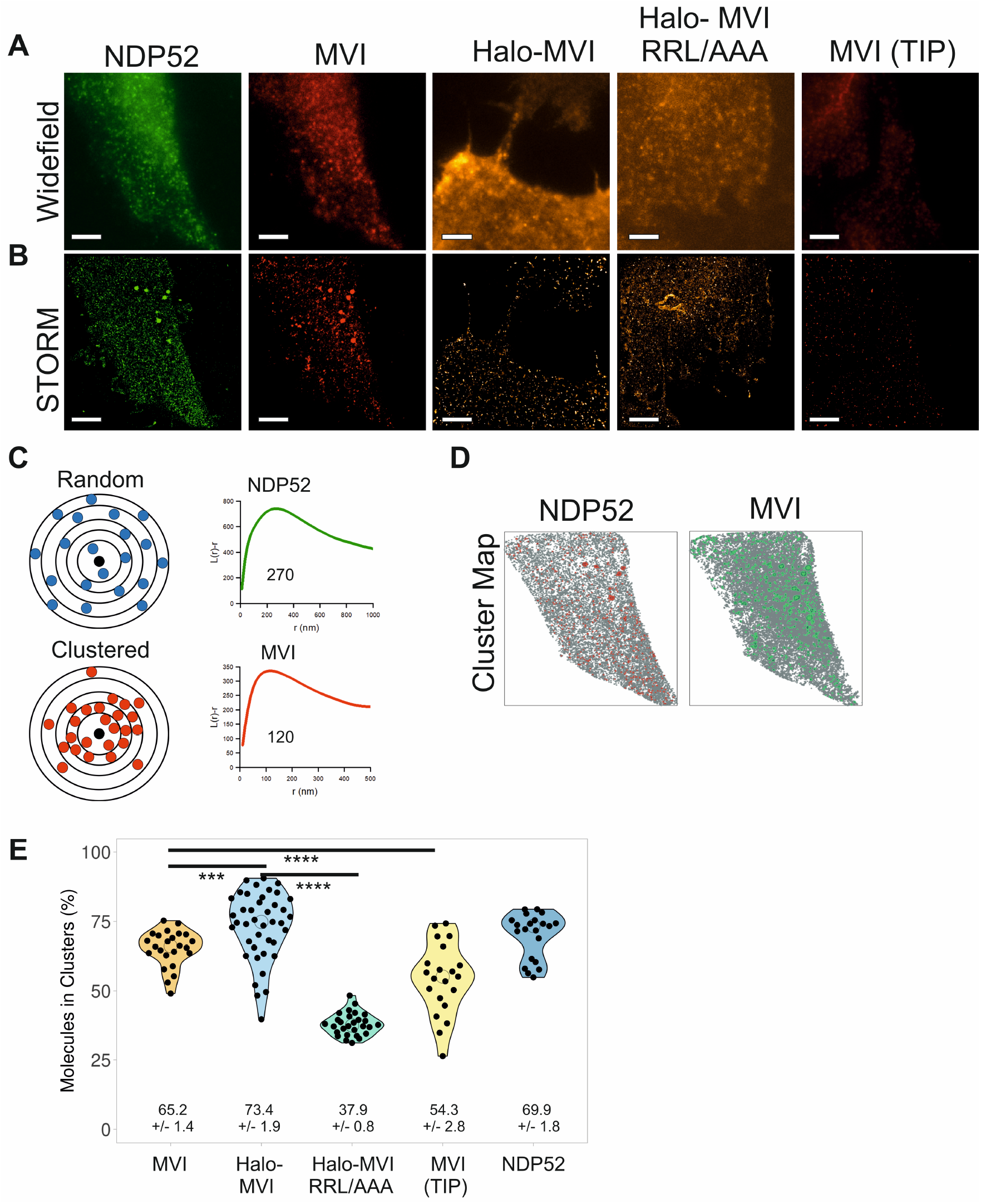
Cellular clustering of myosin VI and NDP52. (A) Widefield Immunofluorescence imaging against endogenous NDP52 (green) and MVI (red) in the presence and absence of TIP, and JF549 staining against transfected Halo-MVI and Halo-MVI(RRL/AAA) in the cytoplasm of HeLa cells (Scale bar 10 μm). (B) STORM render images of the cells shown in (A) (Scale bar 10 μm). Images were acquired as described in the methods. (C) Depiction of a theoretical example of molecular clustering and random distribution. Molecular clustering is assessed by the linearized plot of Ripley’s K function *L(r)-r*, versus *r*, where r is the radius. Distribution can be plotted as an L-function, where a randomly distributed set of molecules is a flat line with a value of zero. The green curve corresponds to the organisation of NDP52 (peak at *r*=270 nm) and the red curve corresponds to the organisation of MVI (peak at *r*=120 nm). (D) Cluster maps based upon the STORM render in (B). Clusters are shown in Red (NDP52) and Green (MVI). (E) Cluster analysis representing the percentage of molecules in a cluster for the conditions in (B). Individual data points correspond to the average number of molecules per cluster in the selected ROI, in an individual cell. The values represent the mean from all the ROIs for each protein (n = >20). (***p <0.001 ****p <0.0001 by two-tailed t-test).

To explore the interaction between MVI and NDP52 within the cell, we performed colocalization analysis of the clusters (Figure 6). Colocalization can be represented by transforming the STORM images (Figure 6A) into colocalization heat maps (Figure 6B), which assigns a different colour to the clusters of each protein depending on the level of colocalization between them. TIP has a clear impact on the fraction of MVI which is colocalised with NDP52, resulting in an almost two-third reduction (Figure 6C). Exploring the cluster data further revealed that, for each protein, the colocalized clusters represent approximately 20 % of all clusters (colocalised and non-colocalised) (Figure 6D). This is not a surprise given that both proteins interact with various other partners and are also involved in distinct pathways. However, the MVI colocalized clusters are 3-fold larger than the non-colocalized ones, and contain 15-fold more molecules. Treatment with TIP reduces the colocalised cluster area by a third and reduces the number of molecules per cluster by 50%. For NDP52, the colocalized clusters are 2-fold larger than the non-colocalized ones, and contain almost 7-fold more molecules.

**Figure 6:**
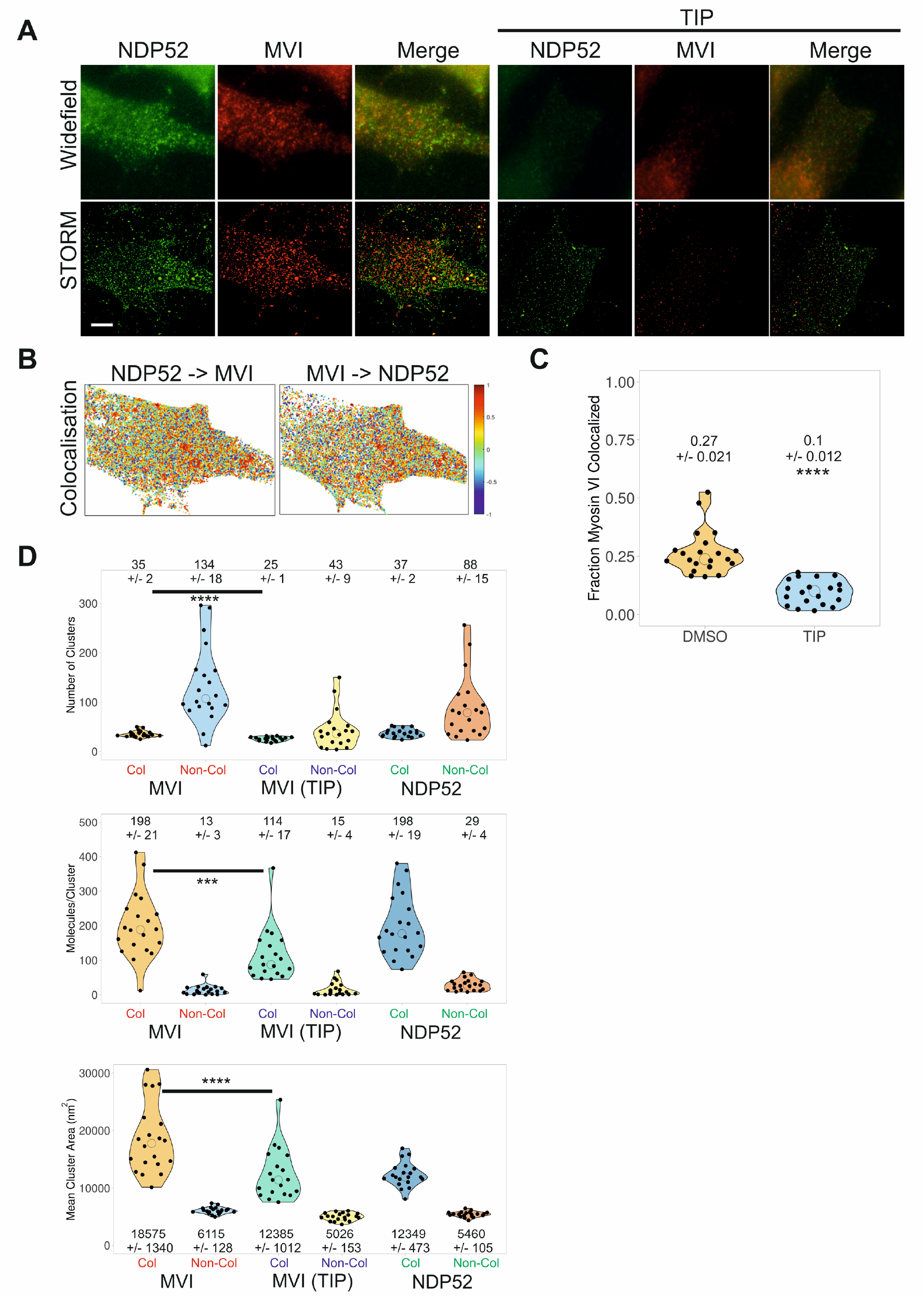
Cluster analysis and colocalization of myosin VI and NDP52. (A) Examples of widefield immunofluorescence imaging against endogenous NDP52 (green) and MVI (red) in the presence and absence of TIP in HeLa cells (Scale bar 10 μm), with their corresponding STORM render. Images were acquired as described in the methods. (B) Cluster colocalization heatmaps corresponding to the STORM renders shown in (A), depicting the colocalisation scores for each molecular cluster for MVI and NDP52. Here, values of 1 (dark red) correspond to perfectly colocalised clusters and −1 (dark blue) to clusters separated from each other. (C) Fraction of MVI colocalised with NDP52 in the presence and absence of TIP. (D) Results of cluster analysis of MVI +/- TIP and NDP52, displaying the distribution and mean values of the number of clusters per cell ROI, number of molecules per cluster and cluster area. The data are broken down into MVI and NDP52 colocalized and non-colocalized clusters. Individual data points correspond to the corresponding average value per cell ROI. The values represent the mean (+/- s.d.) from the ROIs (n = 20) for each condition (****p <0.0001 by two-tailed t-test compared to non-colocalized clusters).

Taken together, the cluster analysis suggests that molecular clustering of MVI does occur within the cell and is promoted by binding partners, while require motor activity. As clustered MVI does not represent freely diffusing molecules, the high local density generated by clustering would enhance the impact of its biochemical properties. Within the confined and denser environment of the clusters, the low affinity interactions of MVI with its partners are more likely to occur. In this way, clustering would ensure that the subsequent unfolding and dimerization of MVI is readily facilitated and implemented, when needed.

### Interactions with binding partners regulate the cellular dynamics of myosin VI

Following the single molecule localisation experiments, we performed live-cell 3D single molecule tracking to observe the impact of binding partners on the dynamics of MVI, using an aberration-corrected multi-focal microscope (acMFM) system [28]. This technique allows the simultaneous acquisition of 9 focal planes covering 4 μm in the *z* axis, with a 20 x 20 μm field of view. We visualised NI isoform of MVI in HeLa using an N-terminal HaloTag fusion [7] labelled with JF549 [29] and then performed single particle tracking.

The trajectories of MVI revealed different populations of molecules, some undergoing confined motion, some random diffusion and others directed movement (Figure 7A). The diffusion constants (*D*) extracted from these trajectories had a mean value of 0.3 μm^2^ s^-1^, under normal conditions, with the large majority of molecules exhibiting slow diffusion (Figure 7B). To better understand the diffusion properties of MVI, we further split these diffusion constants into three groups: a) static/slow moving molecules with D below 0.1 μm^2^ s^-1^, b) molecules with D between 0.1 and 2 μm^2^ s^-1^ and c) fast moving molecules with D above 2 μm^2^ s^-1^ (Figure 7C). We found that MVI molecules are almost equally split between the first two categories, indicating that MVI mainly exists in a static/slow and medium diffusive state. In addition, we assessed the type of particle motion exhibited by MVI by calculating the anomalous diffusion alpha value (Figure 7D). Values below 1 are indicative of confined motion, values above 1 are suggestive of directed motion, whereas values of 1 occur for random diffusion. The alpha value for MVI under normal conditions had a mean value of 0.69, therefore suggesting it mainly undergoes confined movement. This confined motion was further represented using a roseplot (Figure 7E) to depict the angular change in direction within the trajectories. Angles around 180 degrees were enriched and represented trajectories where molecules reversed direction, as would more likely occur for a confined molecule.

**Figure 7:**
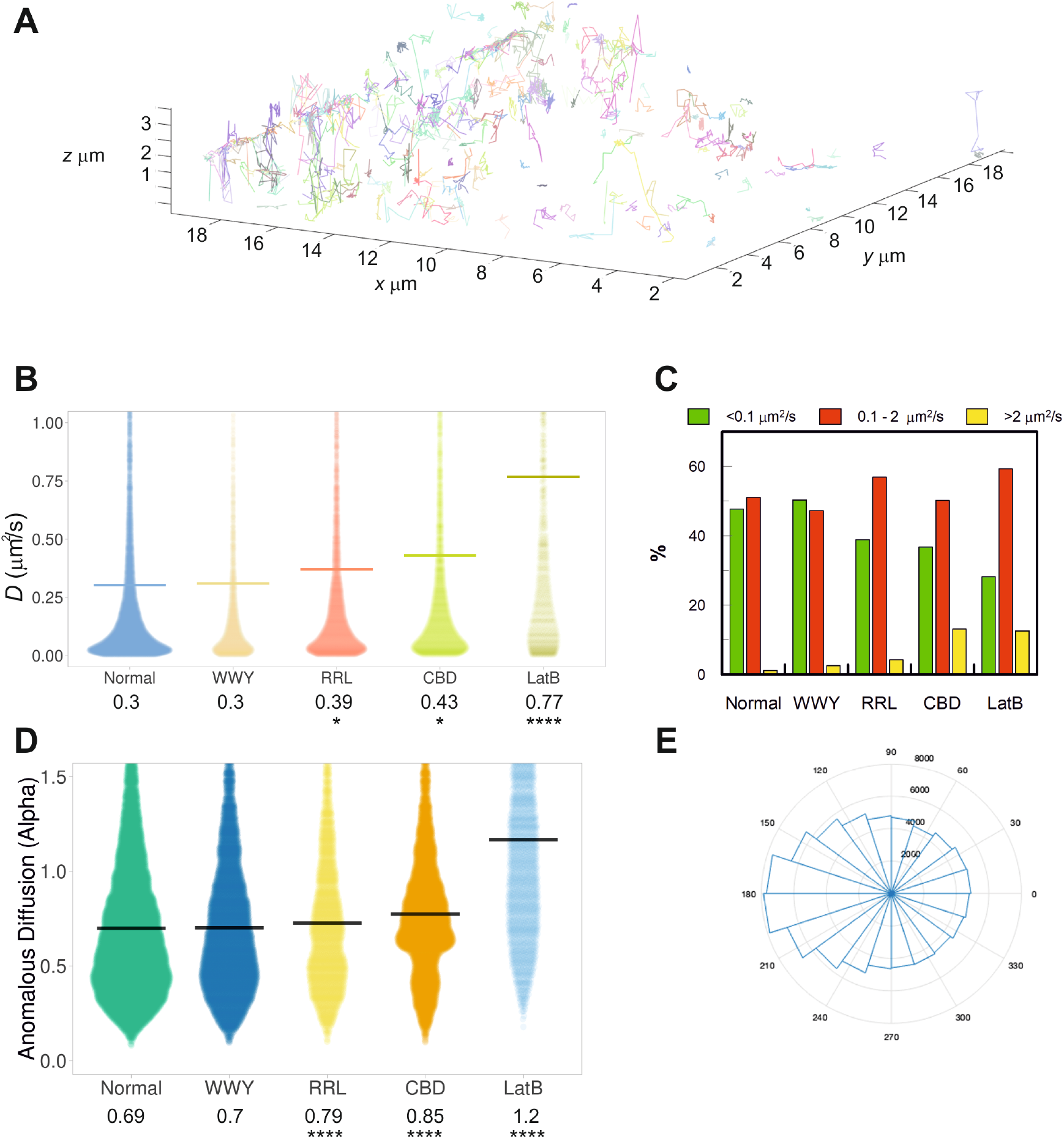
Live cell single-molecule dynamics of myosin VI. (A) Example render of 3D single-molecule trajectories for JF549-labelled Halo-tagged MVI, stably expressed in HeLa cells, under normal conditions. (B) Plot of diffusion constants for wild type (WT) MVI (normal), MVI WWY/WLY (WWY), MVI RRL/AAA (RRL), WT MVI following transient over-expression of GFP-CBD (CBD) and WT MVI following treatment with latrunculin B (LatB). Diffusion constants were derived from fitting trajectories to an anomalous diffusion model, as described in methods. The data represent all trajectories from 100 cells (* p <0.05, ****p <0.0001 by two-tailed t-test compared to normal conditions). (C) Diffusion constants (D) were split into three categories corresponding to static/slow moving (D<0.1 μm^2^ s^-1^), mobile (0.1<D<2 μm^2^ s^-1^) and hyper-mobile fractions (D>2 μm^2^ s^-1^). The percentage of molecules from the experiments in (B) falling into each group were then plotted. (D) Plot of anomalous diffusion alpha values derived from the same experimental set shown in (B), for the indicated conditions. (****p <0.0001 by two-tailed t-test compared to normal conditions). (E) Roseplot representing the angular change in WT MVI diffusion from the trajectories analysed in (B). Angles close to 180° are enriched which suggests molecules are moving backwards and forwards within a space.

To assess the role of binding partners in the cellular dynamics of MVI, we tracked the movement of two MVI mutants in which Dab2 and NDP52 binding was disrupted, namely MVI_WWY/WLY_ and MVI_RRL/AAA_, respectively. MVI_WWY/WLY_ did not show any significant change in its diffusion properties compared to wild type MVI (Figure 7B and C). This is not surprising since, as shown previously, the NI isoform displays selectivity for RRL binding partners, rather than the lower affinity WWY-mediated interactions [6]. Conversely, MVI_RRL/AAA_ led to a significant shift in the distribution of diffusion constants, with a 30 % increase in the mean diffusion constant to a value of 0.39 μm^2^ s^-1^ (Figure 7B). This can also be seen by the decrease in the static/slow moving population (<0.1 μm^2^ s^-1^) and the increase in the medium and highly diffusive pools (0.1 – 2 μm^2^ s^-1^ and >2 μm^2^ s^-1^) (Figure 7C). This was also matched by the significant change in the type of anomalous diffusion exhibited by the MVI mutant molecules, with an increase in the mean anomalous diffusion alpha value to 0.79 (Figure 7D). As evidenced by the dramatic change in the shape of the distribution, disruption of MVI interaction with its RRL binding partners caused a significant shift from confined motion towards random diffusion.

As expected from the biochemical parameters, these data suggest that there is an impact of the binding partners upon the activity of MVI, which influences its cellular dynamics. This was further supported by the effect of transiently over-expressed GFP-tagged CBD into cells stably expressing wild type Halo-MVI. Over-expression of this construct is known to have a dominant negative effect by displacing wild type MVI from binding partner and lipid-based interactions [30]. Indeed, the presence of the CBD led to a considerable increase in the diffusion constant of MVI to a mean value of 0.43 μm^2^ s^-1^ (Figure 7B), an increase in the population of highly diffusive species (Figure 7C), and a shift of the alpha value to mean of 0.85 (Figure 7D). Once again, this is consistent with the STORM measurements which revealed less clustering of MVI RRL mutant. To assess the contribution of actin to the pool of static/slow diffusing molecules, the dynamics of MVI were observed following treatment with an actin polymerization inhibitor, Latrunculin B. Not surprisingly, there was a significant increase in MVI diffusion (Figure 7B and 7C), due to the loss of actin-based interactions.

Overall, our data demonstrate that the interaction of MVI with its binding partners plays an important role in regulating the cellular dynamics of the protein. The increase in random diffusion observed following disruption of these interactions could reflect a loss of interaction with cargo and anchoring sites. It could also reflect loss of direct interactions with actin, given that MVI would then be in a back-folded inactive state.

### Binding partner interactions enhance actin binding

To directly test how binding partners impact actin binding, we performed actin isolation assays with recombinant proteins (Figure 8A/B). As expected, MVI is isolated by Factin. The amount of MVI bound to actin is 50%. The addition of an excess of NDP52 led to above 80% of MVI being bound to actin. As expected, the RRL-AAA mutant did not show the enhanced binding in the presence of NDP52. Whilst these experiments are in alignment with our assays above, this finding contradicts the claims in (Arden 2016) where actin pelleting assays were performed in cell lysates. Based on the findings from Arden et al, we would expect to find these mutant proteins decorating actin filaments. To this end, we transfected HeLa cells with eGFP-WT, RRL-AAA and WWY-WLY MVI (Figure 8C). Some colocalization with actin filaments are observed but there are no obvious differences between mutants and wild type MVI. We therefore conclude we are not enhancing actin binding above wild type levels.

**Figure 8:**
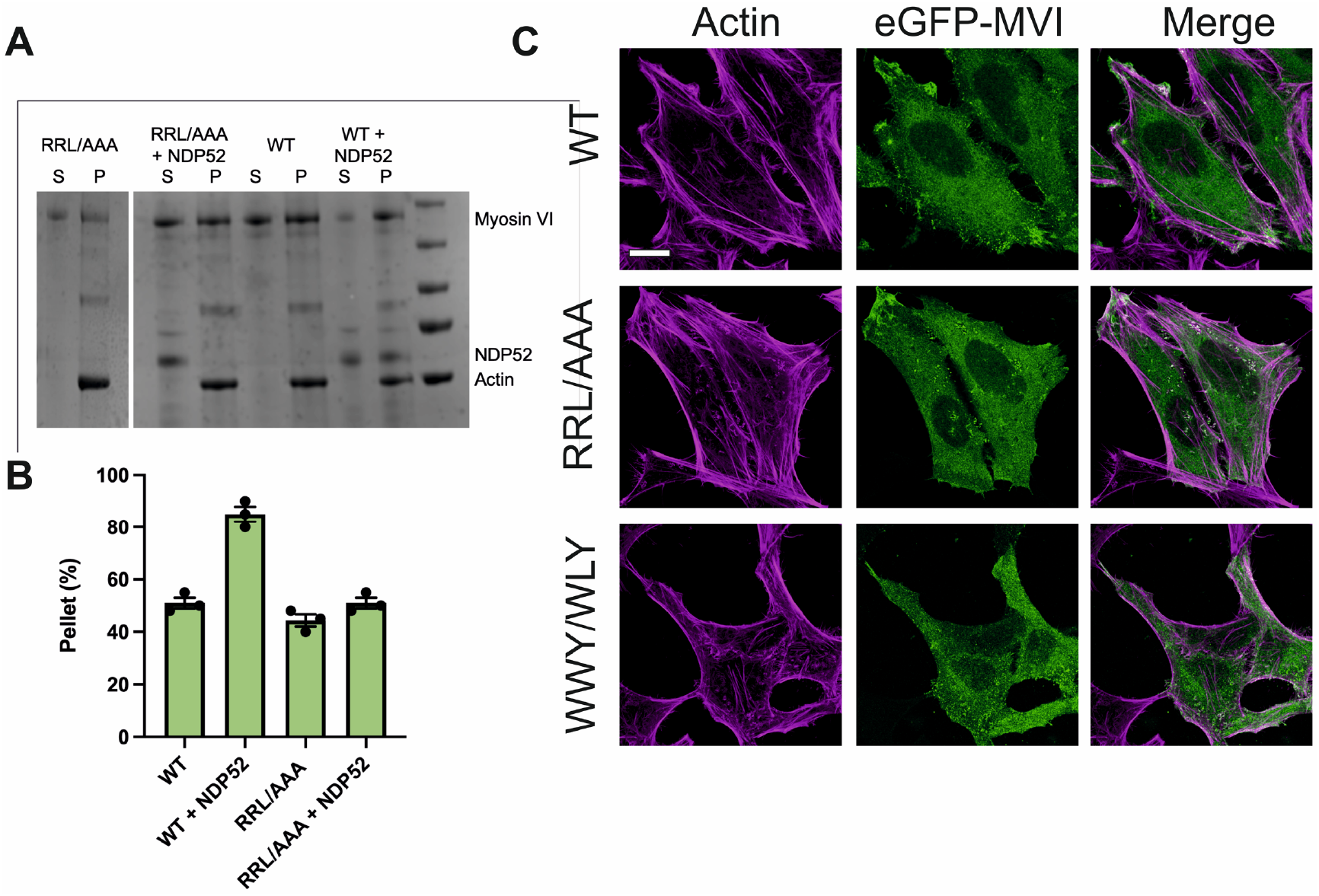
Binding partners enhance actin binding. (A) Actin pull-down of 5 μM myosin VI wild type (WT) or mutant RRL/AAA in the presence and absence of 15 μM NDP52. P and S represent pellet and supernatant, respectively. (B) Quantification of pulldowns in A. (C) Representative images of transiently transfected eGFP-MVI wild type (WT), RRL/AAA and WWY/WLY MVI in HeLa cells combined with phalloidin staining against actin (magenta). Scale bar 10 μm.

## DISCUSSION

This study has provided novel insights into the mechanisms underlying the regulation of MVI and its function within a cell. We have revealed that binding partners directly regulate the structural transitions of MVI from an auto-inhibited backfolded monomer to an active motor, which is linked to actin and cargo (Figure 8A). Our data suggest that this mechanism is generic, irrespective of MVI isoform or choice of binding partners. Importantly, this model does not assume that binding partners must be dimeric. Moreover, pre-steady-state kinetics has allowed us to define that binding partners first bind to the backfolded MVI and subsequently trigger unfolding, rather than binding to a spontaneously unfolded MVI and stabilising that state. This is an important regulatory step which enables binding partners to turn myosin VI on and off.

We have shown that once MVI is unfolded, it has the intrinsic ability to dimerize, as it has been reported before [4, 6, 13, 24, 25, 31]. Here, we have further clarified these models by showing that MVI dimerizes internally and that binding partner dimerization is not the driving factor. We therefore propose that the complex stoichiometry exists as one active binding partner (monomer or dimer) per MVI molecule (Figure 9A). This stoichiometry has been reported from structural studies [25, 32]. Greater differences in oligomerization status may occur between binding partners where monomers or dimers can be favoured. Moreover, larger oligomers, while maintaining 1:1 stoichiometry, could also be produced [32].

**Figure 9:**
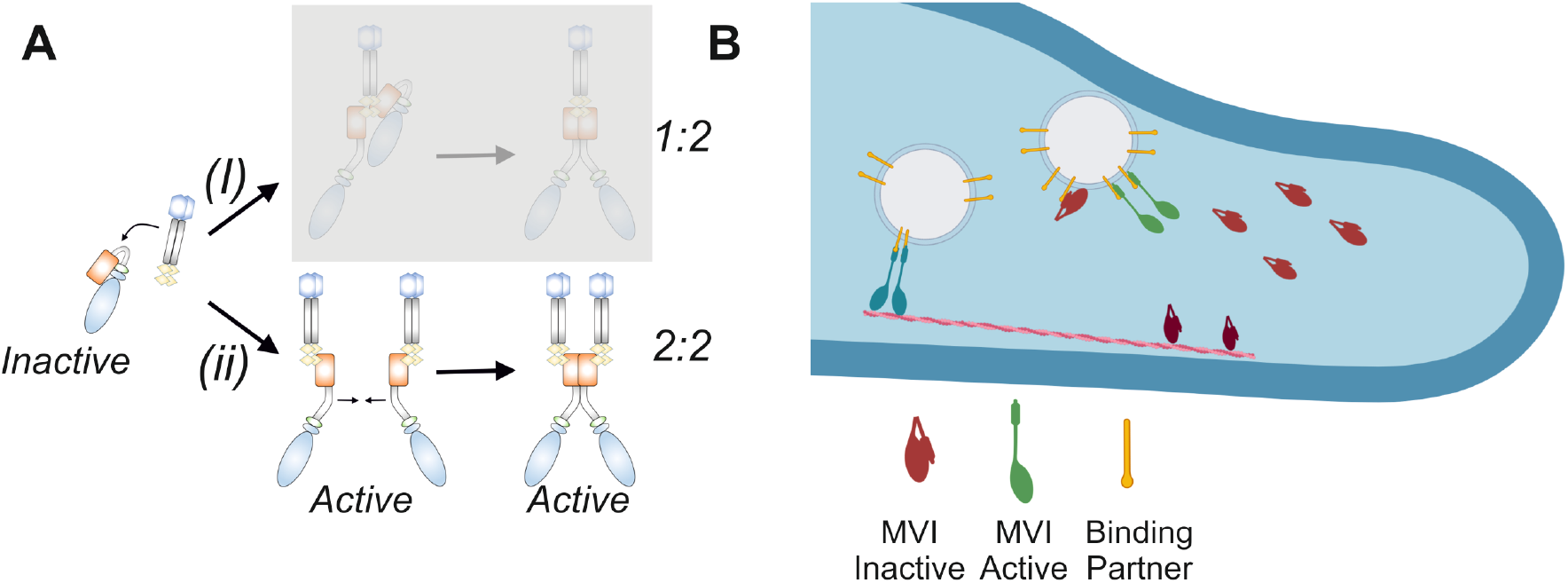
Model describing the activation of myosin VI by binding partners. (A) Two routes of binding partner-dependent dimerization of MVI. (i) Dimeric binding partner (e.g. NDP52) binds to backfolded MVI and triggers its unfolding. A second MVI molecule is then recruited through the binding partner, due to its dimeric nature. This results in a stoichiometry of 1 dimeric partner to 2 MVI motors. This mechanism does not require MVI internal dimerization. (ii) Each MVI molecule is unfolded by an individual binding partner. Unfolding then exposes internal dimerization sequences within MVI, which can result in a dimer complex with a stoichiometry of 2 partners and 2 motors. The data presented here allow option (i) to be excluded. Our data also allows us to propose that the folded state is inactive (non-actin bound) and that the unfolded state (monomer and dimer) becomes active with the ability to bind actin. (B) Recruitment and activation of myosin VI to cellular cargo by binding partners. Monomeric backfolded (inactive) myosin VI (red) can be found freely diffusing in the cytoplasm or bound to actin (dark red). The inactive myosin VI associates with cargo-bound binding partners. This triggers unfolding and activation of the motor in monomeric (green) or dimeric (blue) populations. The active myosin VI can interact with actin filaments and undergo processive movement to facilitate cargo transportation.

Rapid mixing kinetics allowed us to probe the individual mechanistic steps and therefore propose that dimerization can occur rapidly following unfolding of MVI, with unfolding being the rate-limiting step once binding partner interaction occurs.

MVI exists as four alterative spliced isoforms and interacts with a wide range of binding partners. In our study, we assessed the generic nature of MVI unfolding and dimerization, using examples of two MVI isoforms, which represent the two largest structural variants, and two binding partners, which bind at alternative motifs. Although we cannot exclude potential differences in behaviour in other isoforms and with other MVI partners.

Our biochemical analysis has furthered our understanding of how MVI interacts with its binding partners and highlighted the impact of these interactions upon MVI activity. We have previously shown that this relies on the interplay between isoform splicing and the differential affinities between the RRL and WWY binding sites motifs [6]. Here, we have provided insights into the order of events occurring during partner binding, unfolding and dimerization. However, by nature, all these biochemical studies occur under an environment allowing free molecular diffusion, which is strikingly different from the crowded cellular environment. Our high-resolution imaging data shed light on how the biochemical features of MVI measured *in vitro* could translate into the complex cellular context. They revealed that over 60% of MVI molecules are clustered within the mammalian cell. The high local densities within these clusters enhance the probability of interactions with its binding partners, which have been biochemically shown to be in the micromolar range. Moreover, our data have shown that association with binding partners further increases these clusters in size and molecular density. Although this was tested here only for the partner NDP52, it is very likely to apply for all binding partners. The reason is that, within such clusters, multi-valent interactions of MVI to cargo through various binding partners would lead to stable complexes capable of long-range processive movement. Enzymatic clustering has been reported for many cellular processes [33, 34], where is it hypothesised to increase local concentrations to enhance efficiency. Therefore, given the multifunctional and complex nature of MVI intracellular activity, it would not be surprising that this myosin follows a mechanism, which allows stability, efficiency and high level of regulation.

In addition to what was demonstrated biochemically, the impact of binding partners upon MVI is also evident from the live cell single-molecule tracking data. Here, we have shown how mutations, or truncations that lead to the disruption of interactions between MVI and binding partners have a profound effect on the dynamics of the protein inside the cell, leading to loss of the static and slow-moving molecules, which display restrictive diffusion, for more rapidly diffusing molecules exhibiting random motion. To account for such a behavioural switch, we propose that the direct association with the actin cytoskeleton and cellular cargo is decreased when interactions with binding partners are disrupted. In this manner, a backfolded MVI would freely diffuse around the cell and then interact with the cytoplasm when activated by a binding partner on a cargo.

It was previously reported that disruption of MVI – binding partner/cargo interactions leads to an increase in actin binding using a cell-based pull-down assays [35]. However, while imaging was performed with other mutations within their study to show enhanced actin binding, those targeting the RRL and WWY sites were not observed. We performed imaging of these mutations and could not find them decorating actin filaments, as would have been expected based on the study. Instead, the results are consistent with the data presented here, and with our recombinant actin binding assays, where we find that binding partners enhance actin associations.

These observations allowed us to further expand our activation model of MVI. We propose that the back-folded MVI, being inactive, can randomly diffuse in the cell. In this state, MVI can be sequestered by binding partners to cellular cargo or organelles, and get subsequently activated (Figure 9B). Diffusion of MVI in its inactive state is important because it allows sampling of various regions of the cell before recruitment by its partners to function in specific biological processes. Interestingly, we would expect to observe a decrease in ATPase activity if the protein is not bound to actin. However, these changes have not been observed through biochemical analysis, possibly due to the marked differences between the chemical environment of biochemical assays and the cellular environment. Also, within the complex cellular environment, other processes, such as further regulation of the motor by calcium-calmodulin or changes in the phosphorylation status of MVI upon perturbation of binding partner interactions, could lead to inhibition of the ATPase activity, and these would not be observed *in vitro*. These factors remain to be determined.

In summary, our multidisciplinary study provides new insights into the regulation of MVI by binding partners in the cell. The regulation is complex and our knowledge is not complete. It remains unknown as to how monomeric or dimeric states of the protein are selected within the cell, which may be the role of the binding partners to fulfil. Interestingly, due to the internal dimerization of MVI, we suggest that monomeric activities would arise from binding partners activity halting dimerization. Given the established impact of MVI in several diseases, including cancer [19, 36–39] and deafness [35, 40], defining the mechanistic details of its regulation and function is critical for understanding the impact of mutations, truncations, or altered expression of MVI or partners during disease. Overall, these insights provide new avenues for exploring how the activity of this multi-functional motor protein is regulated within the cell and how these processes may be perturbed during disease.

## ACKNOWLEDGEMENTS

We thank the UKRI-MRC (MR/M020606/1) and UKRI-STFC (19130001) for funding. Aberration-corrected multi-focal microscopy was performed in collaboration with the Advanced Imaging Center at Janelia Research Campus, a facility jointly supported by the Howard Hughes Medical Institute and the Gordon and Betty Moore Foundation. We also thank Darren Griffin (University of Kent) and Mike Geeves (University of Kent) for sharing of equipment and reagents, and Satya Khuon (Janelia Research Campus) for assisting with cell culture. The JF549 dye was kindly provided by Luke Lavis (Janelia Research Campus).

## AUTHOR CONTRIBUTIONS

C.P.T. conceived the study. A.dS., N.F., Y.H-G. and C.P.T. designed experiments. N.F. and C.P.T. designed and cloned constructs. A.dS., N.F., Y.H-G., R.E.G. and C.P.T. performed single molecule imaging experiments. Imaging was supported by L.W., M.M-F. and J.A. N.F., R.E.G. and C.P.T. expressed, purified and performed experiments with recombinant proteins. L.W., J.A., T-L.C., A.dS. and C.P.T. contributed to single molecule data analysis. C.P.T. supervised the study. C.P.T. wrote the manuscript with comments from all authors.

## METHODS

### Constructs

A list of constructs and PCR primers are provided in Supplementary Table 1 and 2, respectively. Constructs generated in this work are described below: RRL/AAA and WWY/WLY mutations were made by site-directed mutagenesis using standard Quick-Change site-directed mutagenesis protocol with pLV-Tet0-Halo MVI as the template. All plasmids were verified by DNA sequencing.

### Protein expression and purification in Escherichia coli

Recombinant constructs were expressed in *E.coli* BL21 DE3 cells (Invitrogen) in Luria Bertani media. Proteins were purified by affinity chromatography (HisTrap FF, GE Healthcare). The purest fractions were desalted through a PD10 column (GE Healthcare) to remove imidazole before treatment with TEV protease for 4 hr at 25°C. The samples were then passed through a second HisTrap column. The cleaved protein was further purified through a Superdex 200 16/600 column (GE Healthcare).

### Protein Expression using Baculovirus system

Full-length MVI NI, MVI_1-814_, MVI_1-1060_, MVI_RRL/AAA_ and *Xenopus* calmodulin were expressed in *Sf9* and *Sf21* (*Spodoptera frugiperda*) insect cells using the Bac-to-Bac® Baculovirus Expression System (Invitrogen). *Sf9* cells were cultured in Sf900 media (Gibco). Recombinant bacmids were generated following the manufacturer’s instructions and were transfected into adherent Sf9 cells to generate the P1 viral stock. *Sf9* cells were infected in suspension at 27°C and 100 rpm with 1 in 50 dilution of P1 and P2 viral stocks to yield P2 and P3 stocks, respectively. Finally, expression of recombinant proteins was set up by infecting *sf21* cells with the P3 viral stock in Spodopan media (PAN Biotech). To ensure correct folding of the MVI constructs, cells were simultaneously infected with P3 viral stock of the MVI constructs together with calmodulin at a 0.75 ratio The cells were harvested after 3 days by centrifugation for 15 min at 700xg and at 4 °C and resuspended in ice cold myosin extraction buffer (90 mM KH_2_PO_4_, 60 mM K_2_HPO_4_, 300 mM KCl, pH 6.8), supplemented with Proteoloc protease inhibitor cocktail (Expedeon) and 100 μM PMSF, before proceeding to protein purification. Prior to sonication, an additional 5 mg recombinant calmodulin was added together with 2 mM DTT. After sonication, 5 mM ATP and 10 mM MgCl_2_ were added and the solution was rotated at 4 °C for 30 min before centrifugation (20,000g, 4°C, 30 min). Then, the cell lysate was subjected to the purification. Proteins were purified by affinity chromatography (HisTrap FF, GE Healthcare). The purest fractions were further purified through a Superdex 200 16/600 column (GE Healthcare).

### Protein labelling

Proteins were transferred into 50 mM Na-phosphate (pH 6.5) using a PD10 desalting column. Samples were then incubated with a 5-fold excess of dye for 4 hours, rotating at 4°C. Excess dye was removed using a PD10 desalting column pre-equilibrated with 50 mM Na-Phosphate, 150 mM NaCl and 1 mM DTT. Labelling efficiency was calculated based on the absorbance at 280 nm and the absorbance maximum of the dye. Typical efficiency was 90%, whereby the less than complete labelling was taken as an indicator for a single dye per protein. This was tested for isolated preparations in mass spectroscopy, which revealed both an unlabelled and single labelled population.

### Cell culture and Transfection

HeLa (ECACC 93021013) cells were cultured at 37°C and 5% CO_2_, in Gibco MEM Alpha medium with GlutaMAX (no nucleosides), supplemented with 10% Fetal Bovine Serum (Gibco), 100 units/ml penicillin and 100 μg/ml streptomycin (Gibco). For the transient expression of MVI mutants, HeLa cells grown on glass coverslips were transfected using Lipofectamine 2000 (Invitrogen), following the manufacturer’s instructions. Depending on the construct, 24 h - 72 h after transfection, cells were subjected to further analysis. To inhibit actin polymerization, cells were treated with 1 μM Latrunculin B (Sigma) for 1h at 37 °C.

### Immunofluorescence

HeLa cells were fixed for 15 min at room temperature in 4% (w/v) paraformaldehyde (PFA) in PBS and residual PFA was quenched for 15 min with 50 mM ammonium chloride in PBS. All subsequent steps were performed at room temperature. Cells were permeabilised and simultaneously blocked for 15 min with 0.1 % (v/v) Triton X-100 and 2 % (w/v) BSA in PBS. Cells were then immuno-stained against the endogenous proteins by 1 h incubation with the indicated primary and subsequently the appropriate fluorophore-conjugated secondary antibody (details below), both diluted in 2 % (w/v) BSA in PBS. The following antibodies were used at the indicated dilutions: Rabbit anti-myosin VI (1:200, Atlas-Sigma HPA0354863), Mouse anti-NDP52 (1:250 Abcam ab124372), Donkey anti-mouse Alexa Fluor 488-conjugated (1:250, Abcam Ab181289), Donkey anti-rabbit Alexa Fluor 647-conjugated (1:250, Abcam Ab181347). Coverslips were mounted on microscope slides with Mowiol (10% (w/v) Mowiol 4-88, 25% (w/v) glycerol, 0.2 M Tris-HCl, pH 8.5), supplemented with 2.5% (w/v) of the anti-fading reagent DABCO (Sigma).

### STORM Imaging

Cells were seeded on pre-cleaned No 1.5, 25-mm round glass coverslips, placed in 6-well cell culture dishes. Glass coverslips were cleaned by incubating them for 3 hours, in etch solution, made of 5:1:1 ratio of H_2_O : H_2_O_2_ (50 wt. % in H_2_O, stabilized, Fisher Scientific) : NH_4_OH (ACS reagent, 28-30% NH_3_ basis, Sigma), placed in a 70°C water bath. Cleaned coverslips were repeatedly washed in filtered water and then ethanol, dried and used for cell seeding. Cells were fixed in pre-warmed 4% (w/v) PFA in PBS and residual PFA was quenched for 15 min with 50 mM ammonium chloride in PBS. Immunofluorescence (IF) was performed in filtered sterilised PBS. Cells were permeabilized and simultaneously blocked for 30 min with 3% (w/v) BSA in PBS supplemented with 0.1 % (v/v) Triton X-100. Permeabilized cells were incubated for 1h with the primary antibody and subsequently the appropriate fluorophore-conjugated secondary antibody, at the desired dilution in 3% (w/v) BSA, 0.1% (v/v) Triton X-100 in PBS. The antibody dilutions used were the same as for the normal IF protocol (see above). Following incubation with both primary and secondary antibodies, cells were washed 3 times, for 10 min per wash, with 0.2% (w/v) BSA, 0.05% (v/v) Triton X-100 in PBS or TBS. Cells were further washed in PBS and fixed for a second time with prewarmed 4% (w/v) PFA in PBS for 10 min. Cells were washed in PBS and stored at 4 °C, in the dark, in 0.02% NaN3 in PBS, before proceeding to STORM imaging.

Before imaging, coverslips were assembled into the Attofluor® cell chambers (Invitrogen). Imaging was performed in freshly made STORM buffer consisting of 10 % (w/v) glucose, 10 mM NaCl, 50 mM Tris - pH 8.0, supplemented with 0.1 % (v/v) 2-mercaptoethanol and 0.1 % (v/v) pre-made GLOX solution which was stored at 4 °C for up to a week (5.6 % (w/v) glucose oxidase and 3.4 mg/ml catalase in 50 mM NaCl, 10 mM Tris - pH 8.0). All chemicals were purchased from Sigma.

Imaging was undertaken using the Zeiss Elyra PS.1 system. Illumination was from a HR Diode 642 nm (150 mW) and HR Diode 488 nm (100 mW) lasers where power density on the sample was 7-14 kW/cm^2^ and 7-12 kW/cm^2^, respectively

Imaging was performed under highly inclined and laminated optical (HILO) illumination to reduce the background fluorescence with a 100x NA 1.46 oil immersion objective lens (Zeiss alpha Plan-Apochromat) with a BP 420-480/BP495-550/LP 650 filter. The final image was projected on an Andor iXon EMCCD camera with 25 msec exposure for 20000 frames.

Image processing was performed using the Zeiss Zen software. Where required, two channel images were aligned following a calibration using a calibration using pre-mounted MultiSpec bead sample (Carl Zeiss, 2076-515). The channel alignment was then performed in the Zeiss Zen software using the Affine method to account for lateral, tilting and stretching between the channels. The calibration was performed during each day of measurements.

The images were then processed through our STORM analysis pipeline using the Zen software. Single molecule detection and localisation was performed using a 9 pixel mask with a signal to noise ratio of 6 in the “Peak finder” settings while applying the “Account for overlap” function. This function allows multi-object fitting to localise molecules within a dense environment. Molecules were then localised by fitting to a 2D Gaussian.

The render was then subjected to model-based cross-correlation drift correction. Typical localisation precision was 20 nm for Alexa-Fluor 647 and 30 nm for Alexa-Fluor 488. The final render was then generated at 10 nm/pixel and displayed in Gauss mode where each localisation is presented as a 2D gaussian with a standard deviation based on its precision. The localisation table was exported as a txt for import in to Clus-DoC.

### Clus-DoC

The single molecule positions were exported from Zeiss black version and imported into the Clus-DoC analysis software [27] (https://github.com/PRNicovich/ClusDoC). Cytoplasmic areas were selected as ROIs for cluster analysis. First the Ripley K function was completed on each channel identifying the r max. The r max was then assigned for DBSCAN if one channel was being analysed or Clus-Doc if two channel colcalisation was being analysed. The clustering size was set to a minimum of 5 molecules, with smoothing set at 7 nm and epsilon set at the mean localization precision for the dye. All other analyses parameters remained at default settings. Data concerning each cluster was exported and graphed using Plots of Data.

### Size-exclusion Chromatography and Multi-Angle Light Scattering

100 μl samples of 2mg/ml purified protein, was applied to a Superdex 200 (30 x 1 cm) analytical column (GE Healthcare) equilibrated in 150 mM NaCl, 50 mM Tris.HCl (pH 7.5) and 1 mM DTT and controlled using Waters 626 HPLC at room temperature. Eluted proteins were analysed with Viscotek SEC-MALS 9 and Viscotek RI detector VE3580 (Malvern Panalytical). Molecular mass was determined using OmniSEC software.

### Actin-pelleting Assay

Constructs were incubated at the specified concentrations in reaction buffer (150 mM NaCl, 50 mM Tris.HCl (pH 7.5), 1 mM MgCl_2_ and 1 mM DTT), for 10 min at RT. 5 μM F-actin (mixture 20% Biotinylated actin (Cytoskeleton Inc.)) was added to the mixture and incubated at RT for 10 min. Streptavidin Dynabeads (M-280) (Invitrogen) were washed 3 times according to the manufacturer’s instructions and then finally in reaction buffer. 0.2 mg/ml beads were added to the samples. The samples were incubated for 5 min at RT before magnetic isolation. The isolated sample was resuspended in an equal volume to the supernatant. The samples were analyzed by SDS-PAGE.

### Multi-focal Imaging and Particle Tracking Analysis

Cells stably or transiently expressing Halo-tag constructs were labelled for 15 min with 10 nM HaloTag-JF549 ligand, in cell culture medium at 37°C, 5% CO_2_. Cells were washed for 3 times with warm cell culture medium and then incubated for further 30 min at 37°C, 5% CO_2_. Cells were then washed three times in pre-warmed FluoroBrite DMEM imaging medium (ThermoFisher Scientific), before proceeding to imaging.

Single molecule imaging was performed using an aberration-corrected multifocal microscope (acMFM), as described by Abrahamsson et al. [28]. Briefly, samples were imaged using 561nm laser excitation, with typical irradiance of 4-6 kW/cm^2^ at the back aperture of a Nikon 100x 1.4 NA objective. Images were relayed through a custom optical system appended to the detection path of a Nikon Ti microscope with focus stabilization. The acMFM detection path includes a diffractive multifocal grating in a conjugate pupil plane, a chromatic correction grating to reverse the effects of spectral dispersion, and a nine-faceted prism, followed by a final imaging lens.

The acMFM produces nine simultaneous, separated images, each representing successive focal planes in the sample, with ca. 20 μm field of view and nominal axial separation of ca. 400nm between them. The nine-image array is digitized via an electron multiplying charge coupled device (EMCCD) camera (iXon Du897, Andor) at up to 32ms temporal resolution, with typical durations of 30 seconds.

3D+t images of single molecules were reconstructed via a calibration procedure, implemented in Matlab (MathWorks), that calculates and accounts for (1) the interplane spacing, (2) affine transformation to correctly align each focal plane in the xy plane with respect to each other, and (3) slight variations in detection efficiency in each plane, typically less than ±5-15% from the mean.

Reconstructed data were then subject to pre-processing, including background subtraction, mild deconvolution (3-5 Richardson-Lucy iterations), and/or Gaussian denoising prior to 3D particle tracking using the MOSAIC software suite [41]. Parameters were set where maximum particle displacement was 400 nm and a minimum of 10 frames was required. Tracks were reconstructed, and diffusion constants were extracted via MSD analysis [42] using custom Matlab software assuming an anomalous diffusion model.

### Steady-state ATPase Activity of MVI

Ca^2+^-actin monomers were converted to Mg^2+^-actin with 0.2 mM EGTA and 50 μM MgCl_2_ before polymerizing by dialysis into 20 mM Tris.HCl (pH7.5), 20 mM imidazole (pH 7.4), 25 mM NaCl and 1 mM DTT. A 1.1 molar equivalent of phalloidin (Sigma) was used to stabilize actin filaments, as previously described [43].

Steady-state ATPase activities were measured at 25 °C in KMg50 buffer (50 mM KCl, 1 mM MgCl_2_, 1 mM EGTA, 1 mM DTT, and 10 mM imidazole, pH 7.0). Supplemented with the NADH-coupled assay components, 0.2 mM NADH, 2 mM phosphoenolpyruvate, 3.3 U ml^-1^ lactate dehydrogenase, 2.3 U ml^-1^ pyruvate kinase and various actin concentrations (0 – 30 μM). The final [Mg.ATP] was 5 mM and MVI concentration was 100–300 nM. The assay was started by the addition of MVI. The change in absorption at OD_340_ nm was followed for 5 min. The *k*_cat_ and *K*_actin_ values were determined by fitting the data to equation 1.

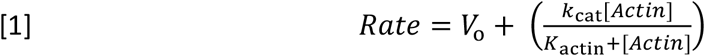

*V*_o_ is the basal ATPase activity of MVI, *k*_cat_ is the maximum actin-activated ATPase rate and *K*_actin_ is the concentration of actin needed to reach half maximal ATPase activity.

### Titration measurements

All reactions were performed at 25 °C in a buffer containing 50 mM Tris·HCl (pH 7.5), 150 mM sodium chloride and 1 mM DTT in a final volume of 100 μL. Measurements were performed using a ClarioStar Plate Reader (BMG Labtech). Intensity measurements were performed at the following wavelengths: FITC (ex. 490nm), Alexa Fluor 555 (ex. 555nm). FITC to Alexa Fluor 555 FRET measurements were performed using the following wavelengths ex. 470nm and em. 575nm.

### Stopped flow measurements

A HiTech SF61DX2 apparatus (TgK Scientific Ltd, Bradford-on-Avon, UK) with a mercury-xenon light source and HiTech Kinetic Studio 2 software was used [44, 45]. For FRET experiments, excitation was at 495 nm with emission through a 570 nm cutoff filter (Schott Glass). For Cy3B, excitation was at 550 nm with emission through a 570 nm cut-filter (Schott Glass). In all experiments, the quoted concentrations are those in the mixing chamber, except when stated. All experiments were performed at 25°C in 50 mM Tris-HCl, 150 mM NaCl, 1 mM DTT and 3 mM MgCl_2_. The dead time of the stopped-flow instrument was ~2 ms: during this initial time no change in fluorescence can be observed.

### Analysis of kinetic data

For the FRET titrations: The 575 nm intensity data was corrected for the increase in intensity due to a small direct excitation. This background signal was subtracted from the dataset to leave the FRET values. The titration curves for the MVI_TAIL_ interactions were fitting to a binding quadratic equation, Equation 2:

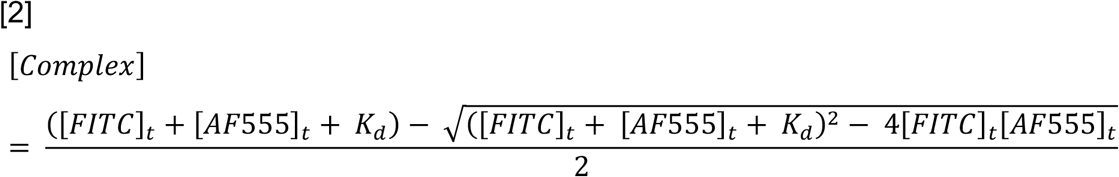

### Graphics

Unless stated, data fitting and plotting was performed using Plots of data [46] and Grafit Version 5 (Erithacus Software Ltd). Cartoons were generated using the BioRender software.

### Data Availability

The data supporting the findings of this study are available from the corresponding author on request.

