## Supplementary Data for "Binding partners regulate unfolding of myosin VI to activate the molecular motor"

**Supplementary Table 1. Recombinant DNA.**

| **Construct (Residue numbers)** | **Source** |
| --- | --- |
| Xenopus pFastBac1 Calmodulin (1-end) | J. Sellers (NIH) |
| Human pET151 NDP52 (1-end) | Ref Fili et al 2017 |
| Human pET151 EGFP-MVI_TAIL(NI)_-RFP (814-1253) | Ref Fili et al 2017 |
| Human pET151 EGFP-MVI_TAIL(LI)_-RFP (814-1284) | Synthetic Gene – This study |
| Human pET151 MVI_TAIL_ 814-1253 | Ref Fili et al 2017 |
| Human pET151 MVI_TAIL_ 814-1060(NI) | Ref Fili et al 2017 |
| Human pET151 MVI_TAIL_ 814-1091(LI) | Synthetic Gene – This study |
| Human pET28 CBD 1060-1253 | Ref Fili et al 2017 |
| Human pFastbacHTB NI MVI (1-1253) | Ref Fili et al 2017 |
| Human pFastbacHTB MVI (1-814) | Ref Fili et al 2017 |
| Human pFastbacHTB MVI (1-1060) | Ref Fili et al 2017 |
| Xenopus pET28 Cys Calmodulin | Ref Fili et al 2017 |
| Human pET151 Dab2 (649-770) | Ref Fili et al 2020 |
| Human pEGFP-C3 CBD 1060-1253 | F. Buss (CIMR) |
| Human pLV-Tet0-Halo-MVI RRL/AAA | This study |
| Human pLV-Tet0-Halo-MVI WWY/WLY | This study |
| Human pEGFP-C3 NI MVI | Ref Fili et al 2017 |
| Human pEGFP-C3 NI MVI RRL/AAA | Ref Fili et al 2020 |
| Human pEGFP-C3 NI MVI WWY/WLY | Ref Fili et al 2020 |
| Human pcDNA3.1 Halo-MVI | Synthetic Gene – This study |

**Supplementary Table 2. PCR Primers.**

| **Sequence** | **Use** |
| --- | --- |
| CTTGCAGAGAAGAATTTCATGCGGCAGCAAAAGTGTATCATGCTTGGAAATC | RRL/AAA For |
| GATTTCCAAGCATGATACACTTTTGCTGCCGCATGAAATTCTTCTCTGCAAG | RRL/AAA Rev |
| CCCTCAGAGTAAGAAAAAAAGGCTGGTTGTATGCCCATTTTGATGGACC | WWY/WLY For |
| GGTCCATCAAAATGGGCATACAACCAGCCTTTTTTCTTACTCTGAGGG | WWY/WLY Rev |

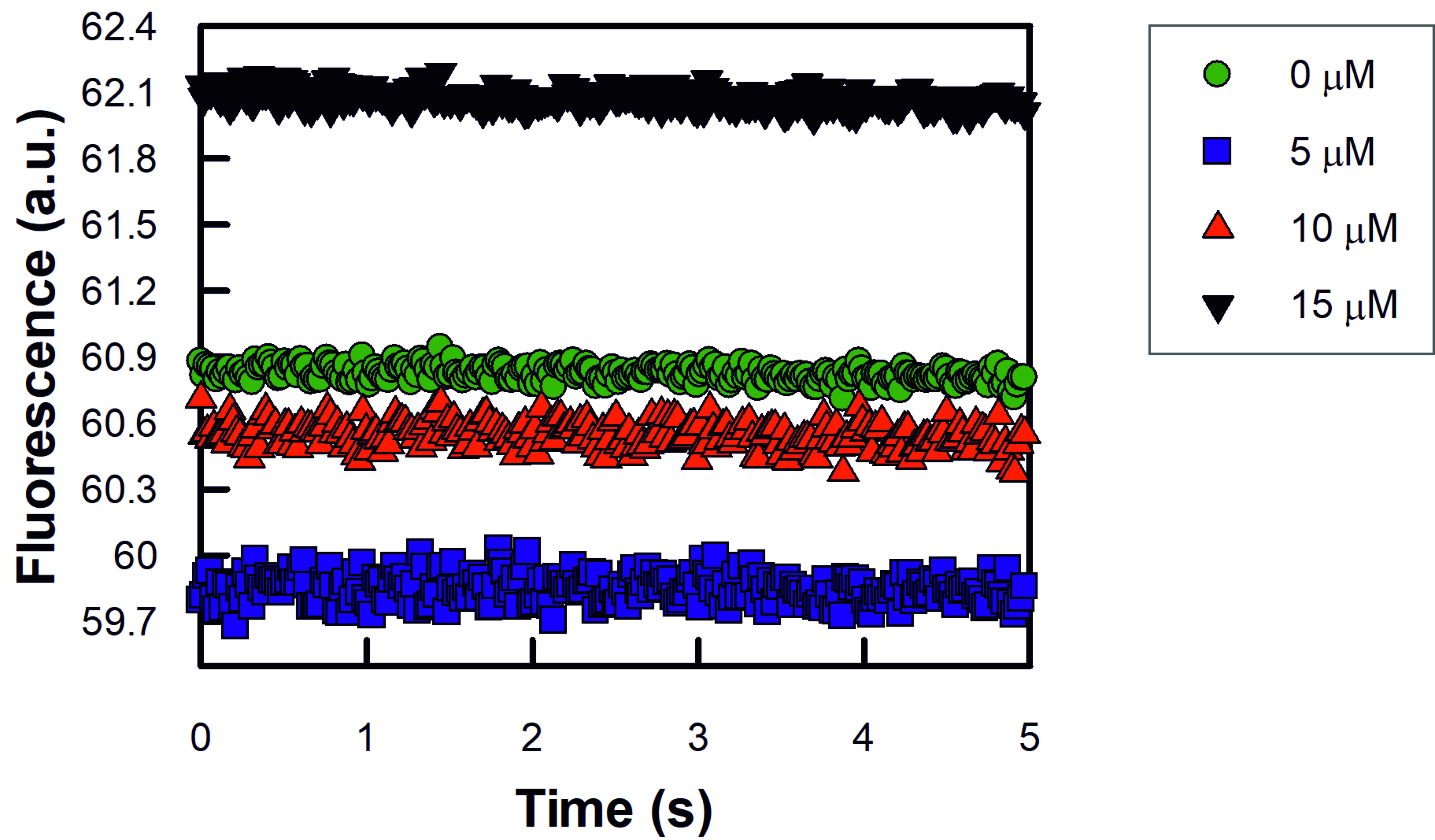

**Supplementary Figure 1.** Representative stopped-flow fluorescence traces following rapid mixing of 1 μM Cy3B-calmodulin and unlabelled NDP52 at the stated concentrations. Experiments were performed as described in the methods.

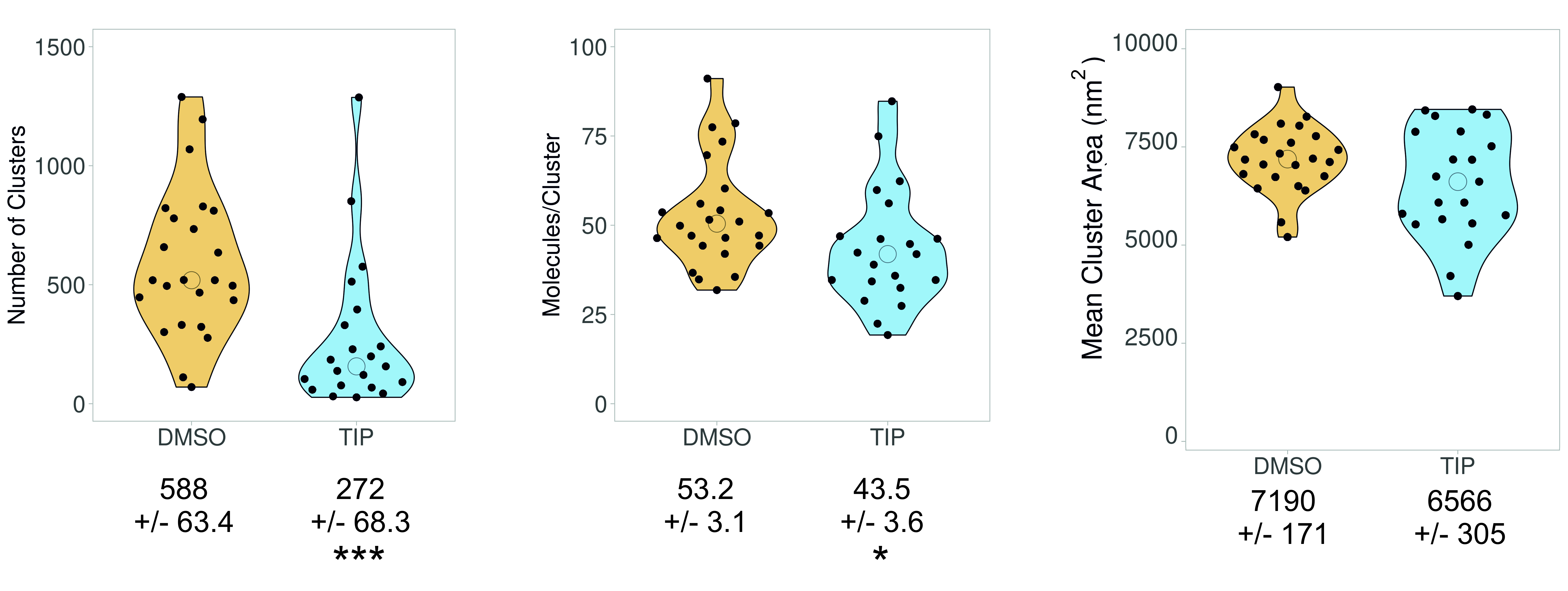

**Supplementary Figure 2.** Cluster analysis of MVI in the presence and absence of TIP, as described in Figure 5. Individual data points correspond to the average number of molecules per cluster in the selected ROI, in an individual cell. The values represent the mean from all the ROIs for each protein (n = >20). (*p <0.05 ***p <0.001 by two-tailed t-test).
